# hERG-LTN: A New Paradigm in hERG Cardiotoxicity Assessment Using Neuro-Symbolic and Generative AI Embedding (MegaMolBART, Llama3.2, Gemini, DeepSeek) Approach

**DOI:** 10.1101/2025.02.17.638731

**Authors:** Delower Hossain, Fuad Al Abir, Jake Y Chen

## Abstract

Assessing adverse drug reactions (ADRs) during drug development is essential for ensuring the safety of new compounds. The blockade of the Ether-a-go-go-related gene (hERG) channel plays a critical role in cardiac repolarization. Computational predictions of hERG inhibition can help foresee drug safety, but current data-driven approaches have limitations. Therefore, a new paradigm that bridges the gap between data and knowledge offers an alternative for advancing precision pharmacogenomics in assessing hERG cardiotoxicity. This study aims to develop a reasoning-based, in silico, robust model for predicting drug-induced hERG inhibition, facilitating new drug development by reducing time and cost, supporting downstream in vitro and in vivo testing. In this study, we constructed a new cohort, UnihERG_DB, by sourcing data from ChEMBL, PubChem, BindingDB, GTP, hERG Karim’s, and hERG Blocker’s bioactivity databases. The final dataset comprises 20,409 structures represented as SMILES (Simplified Molecular Input Line Entry System), labeled as hERG blockers (IC50 < 10 µM) or non-hERG blockers (IC50 ≥ 10 µM). Molecular features were extracted using Morgan and CDK fingerprints. Furthermore, we explored embedding feature computation using cutting-edge Large Language Models, including NVIDIA MegaMolBART, LLaMA 3.2, Gemini, and DeepSeek. Finally, we utilized the Logic Tensor Network (LTN), an advanced AI framework, to train and develop the hERG predictive model. Model performance was evaluated using two benchmarks: External Test-1 and hERG-70. The Logic Tensor Network (LTN) outperformed several models, including CardioTox, M-PNN, DeepHIT, CardPred, OCHEM Predictor-II, Pred-hERG 4.2, Random Forest, and Gradient Boosting. On the External Test-1 dataset, LTN achieved an accuracy of 0.931, a specificity of 0.928, and a sensitivity of 0.933. Furthermore, on the hERG-70 benchmark, LTN achieved an accuracy (ACC) of 0.827, a specificity (SPE) of 0.890, and a correct classification rate (CCR) of 0.833. Overall, the Neuro-Symbolic AI approach sets a new standard for hERG-related cardiotoxicity assessment, yielding competitive results with current state-of-the-art (SOTA) models, and highlights its potential for advancing precision pharmacogenomics in drug discovery and development ([GitHub](https://github.com/hossain013/hERG-LTN.git)).

## I. Introduction

The study of Cardiotoxicity and its impact on drug discovery has become a spirited area of research in recent years. The human Ether-à-go-go-Related Gene (hERG) encodes a potassium channel essential for cardiac repolarization. Drugs that block this channel can lead to prolonged Q.T. intervals on an electrocardiogram (ECG), increasing the risk of arrhythmias as well as sudden cardiac death. Therefore, accurately classifying this type of Adverse Drug Reactions (ADRs) is pivotal during drug development. Several studies [[1]-[3] have leveraged A.I. methodologies to assess Drug-Induced Cardiotoxicity (DICT). Our studies have identified various QSAR (Quantitative Structure-Activity Relationship) models [4]-[5] that employ machine and deep learning strategies [6]-[20], including auto-QSAR modeling, to predict hERG inhibition. Current state-of-the-art models include CToxPred-hERG [21], CardioTox [22], M-PNN [1], DeepHIT [24], CardPred [25], ADMETlab 2.0 [26], ADMETsar 2.0 [27], OCHEM Predictor-I and II [28], Pred-hERG 4.2 [1], and CardioGenAI [30].

Unlike existing predictive approaches that rely solely on typical molecular properties or conventional A.I. experiments, methods of integrating knowledge with data are essential in healthcare, where reasoning, explainability, and interpretability is paramount. However, this integration has been inadequately addressed in hERG inhibition prediction research. In recent years, Neuro-symbolic (NeSy) approaches [31], a new dimension of AI, have garnered significant attention for combining neural networks’ strengths with symbolic reasoning, resulting in more interpretable and insightful predictions. Our study identified implemented several innovative NeSy models across both healthcare and non-healthcare domains, including (Gene Sequence) KBANN [32], (Diabetic Retinopathy) ExplainDR [33], (Link Prediction) NeuralLP [34], (Ontology) RRN [35], NSRL [36], Neuro-Fuzzy [37], FSKBANN [38], DeepMiRGO [39], NS-VQA [40], DFOL-VQA [41], LNN [42], NofM [43], PP-DKL [44], FSD [45], CORGI [46], NeurASP [47], XNMs [48], Semantic Loss [49], NS-CL [50], and LTN [51]. Moreover, the emergence of generative AI is poised to revolutionize the precision pharmacogenomics industry by accelerating drug discovery, particularly in chemical compound research.

The aim of this study was to investigate data- and knowledge-driven approaches (Neuro-Symbolic AI) by extracting features from Large Language Models (LLMs) to enhance in silico cardiac predictive modeling. We developed a large, diversified cohort by procuring data from ChEMBL, PubChem, BindingDB, GTP, hERG Karim’s, and hERG Blocker’s bioactivity databases, surpassing the scope of existing research. By conducting diverse experiments, we compared the performance of molecular fingerprints, descriptors, and embeddings generated by NVIDIA’s MegaMolBART, LLaMA 3.2, Gemini, and DeepSeek-QWEN_1.5b with the novel AI approach, Logic Tensor Network (LTN), to provide valuable insights into hERG-related cardiotoxicity prediction. This new mechanism not only outperformed existing state-of-the-art models on the External Test-1 dataset but also demonstrated significant improvements in the hERG-70 benchmark, setting a new standard in the field of cardiotoxicity assessment.

## II. Materials and Methods

This section outlines the procedures employed in experiments to assess the performance of our proposed hERG-LTN approach. The hERG-LTN paradigm leverages several features engineering strategies, including chemical LLM embeddings generated by the MegaMolBART, LLaMA 3.2, Gemini, and DeepSeek-QWEN_1.5b model. This section covers the entire pipeline, including materials, data preprocessing, feature extraction, network architecture, LTN knowledge-based settings, the training and inference phases, and the system of measurements used to evaluate the model’s performance against state-of-the-art models.

### A. Data acquisition

In this study, a new cohort was constructed, UnihERG_DB, by sourcing data from several bioactivity databases: ChEMBL [53], hERG Karim’s (TDC), BindingDB [54], PubChem [55], Guide to Pharmacology (GTP)[56], and hERG Blockers [57]. The ChEMBL data was obtained using ID CHEMBL240, BindingDB via the link, PubChem in CSV format with NCBI Gene 3757 (link), and GTP via the corresponding link. The hERG Karim’s and hERG Blocker’s datasets were collected from the TDC website using the Python package PyTDC. After curation, the UnihERG_DB database consists of 20,409 trainable compounds (Table 1). The final dataset comprises 20,409 structures represented as SMILES (Simplified Molecular Input Line Entry System). labeled as hERG blockers (IC50 < 10 µM) or non-hERG blockers (IC50 ≥ 10 µM).

**Table 1:** UnihERG_DB. Data Distributions

| Source | Blocks | Non-blocks | Both |
| --- | --- | --- | --- |
| ChEMBL | 5460 | 6320 | 11780 |
| hERG Karim | 2628 | 2469 | 5097 |
| BindingDB | 952 | 955 | 1907 |
| PubChem | 719 | 749 | 1468 |
| hERG Blockers | 65 | 88 | 153 |
| GTP | 2 | 2 | 4 |
| **Total** | **9826** | **10583** | **20409** |

### B. Data Preprocessing

Both hERG Karim and hERG Blocker’s datasets offered preprocessed molecular data and corresponding binary IC50 information representing blockers (IC50<10 μM) and non-blockers (IC50>10 μM). However, numerical IC50 measurements in nM were given in ChEMBL, BindingDB, and GTP, but those in μM were given in PubChem. We harmonized all units into Micromolar (μM), followed by thresholding as blockers (IC50 <10 μM) or non-blockers (IC50>10 μM). We used MolVS Python-Package to standardize SMILES and applied a standard scaler by scikit-learn for embedding features. Finally, we dropped duplicates and NaN values.

### C. Feature extraction

PadelPy is a widely used tool for converting SMILES strings into numerical descriptors, offering more than 10 types of descriptors, including CDK, CDKextended, AtomPairs2DCount, AtomPairs2D, EState, CDKgraphonly, KlekotaRothCount, KlekotaRoth, MACCS, SubstructureCount, and Substructure. Each descriptor type provides different features, with CDK (1024 features) consistently delivering superior results. In this study, we extracted descriptors, fingerprints, and embedding. Using CDK fingerprints, additionally Morgan descriptors (nBits= 2048 and radius = 2) via RDKit. Finally, embeddings were generated by NVIDIA’s advanced LLM model, MegaMolBART [52], by following the instructions on GitHub to configure the necessary environment in Docker. In addition, we computed LLaMA 3.2, Gemini, and DeepSeek-QWEN_1.5b embeddings as well.

### D. Model Architecture / Development

The LTN [51] framework was employed to build the hERG-LTN classifier. The merit of this approach is its capability to tackle the limitations of traditional deep learning systems, which often struggle with tasks requiring reasoning, interpretability, with symbolic manipulation, and knowledge integration. The LTN architecture consists of two key components: a logic component and a neural network. The visual architecture of the classification model is shown in Figure 1. The logical mechanism includes a set of axioms defined based on domains (features and labels), variables, constants (classes), and predicates (p), as detailed in Table 2. During backpropagation, weights are updated using LTN loss functions (Equation 1). We constructed the hERG-LTN classifier by building a Multi-Layer Perceptron (MLP) with four layers, input units for CDK (1024), Morgan (2048), and various types of LLM embeddings features for the hRG-LTN training phase. The hidden layers consist of 16, 8, and 2 units, with Elu activation, a batch size of 64, an Adam optimizer with a learning rate of 0.001, and a seed of 42 with 500 epochs. The LTN knowledge-based setup is detailed in Table 2. In the training phase, the whole dataset was split by scaffolding into 90% for training, 10% for validation (used for UnihERG_DB Dataset). This initial setup provided baseline results. Subsequently, we trained the model on the significant amount dataset and evaluated it using external test-1 and the hERG-70 benchmark.

**Table 2:** LTN Knowledge-based Setting for hERG Classification.

| Table 2: LTN Knowledge-based Setting for hERG Classification |  |
| --- | --- |
| Contents | Classification |
| Define Axioms | $\forall x_A, p(x_A, l_A)$: all the examples of class $A$ ($non - blocks = 0$) should have a label $l_A$<br>$\forall x_B, p(x_B, l_B)$: all the examples of class $B$ ($blocks = 1$) should have a label $l_B$ |
| Axioms (rules, knowledge base) | $\mathcal{K} = \forall x_A p(x_A, l_A), \forall x_B p(x_B, l_B)$ |
| SatAgg is given by | $SatAgg_{\phi \in \mathcal{K}} \mathcal{G}_{\theta, x \leftarrow D}(\phi)$ |
| Learning & Loss | $L = \left( 1 - SatAgg_{\phi \in \mathcal{K}} \mathcal{G}_{\theta, x \leftarrow B}(\phi) \right)$ |
| **Note:** This table was developed inspired by the official LTN |  |

**Fig. 1:**
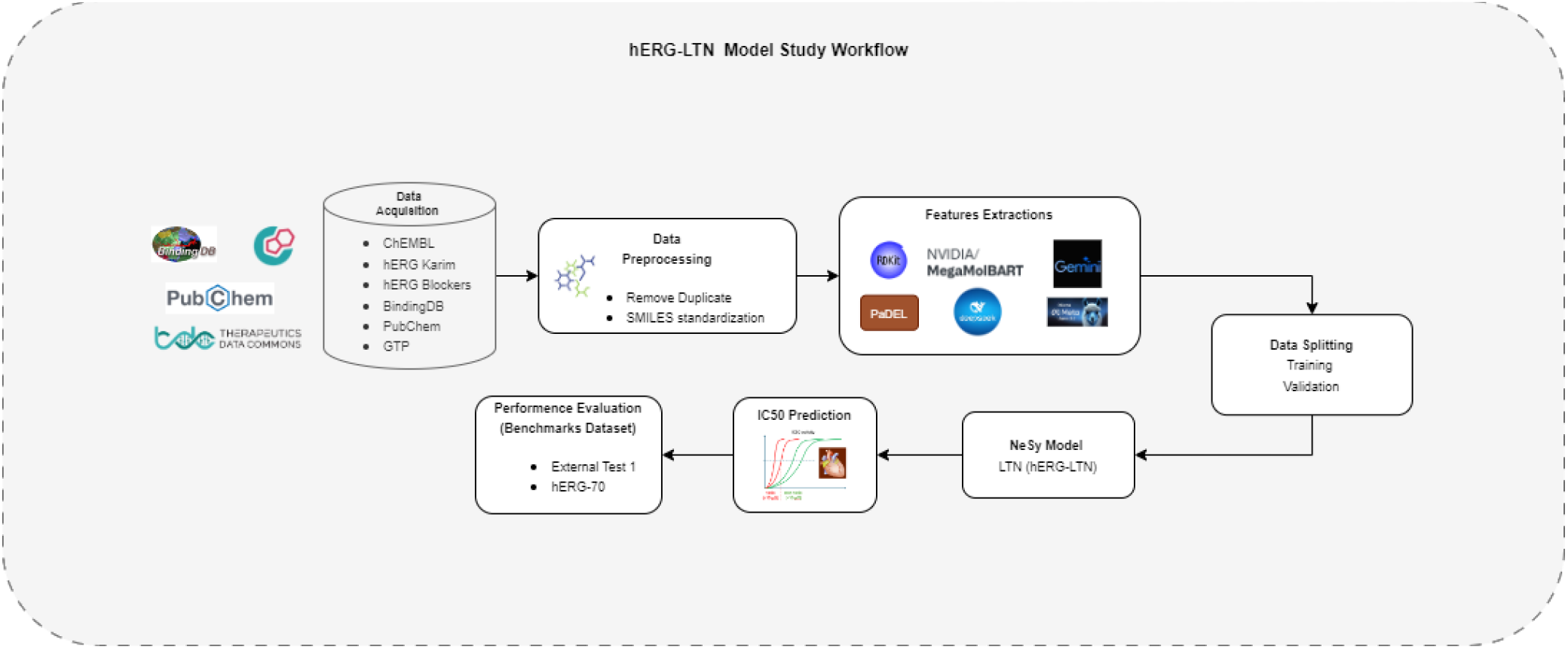
Study work flow of hERG-LTN Model

Here,

The pMeanError aggregator

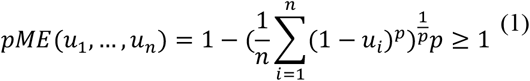

- SatAgg: This stands for “Satisfaction Aggregator”
- ϕ∈K: This part indicates that ϕ (phi) belongs to the set K. ϕ is often used to represent a predicate.
- 𝒢(θ): This is denoted by grounding (𝒢) with parameters θ. θ represents a set of parameters or weights in a model.
- x←D: *D* the data set of all examples (domain).
- The input to the functions SatAgg and 𝒢(θ)

## III. Results

The performance of the proposed hERG-LTN approach will be described in this section, which was trained using various molecular representations for hERG Cardiotoxicity prediction. The hERG-LTN approach utilizes Fingerprint (CDK), Descriptors (Morgan), and LLM-generated embeddings, and its performance is compared to existing state-of-the-art models. The main goal is to emphasize the advantages of the hERG-LTN model, which mainly incorporates logical reasoning for enhanced decision-making in cardiac drug-induced. Please be informed that Table 3 and Table 4 are sort of ablation study across different features input while Table 5, and Table 6 refers to benchmarking comparison.

**Table 3:**
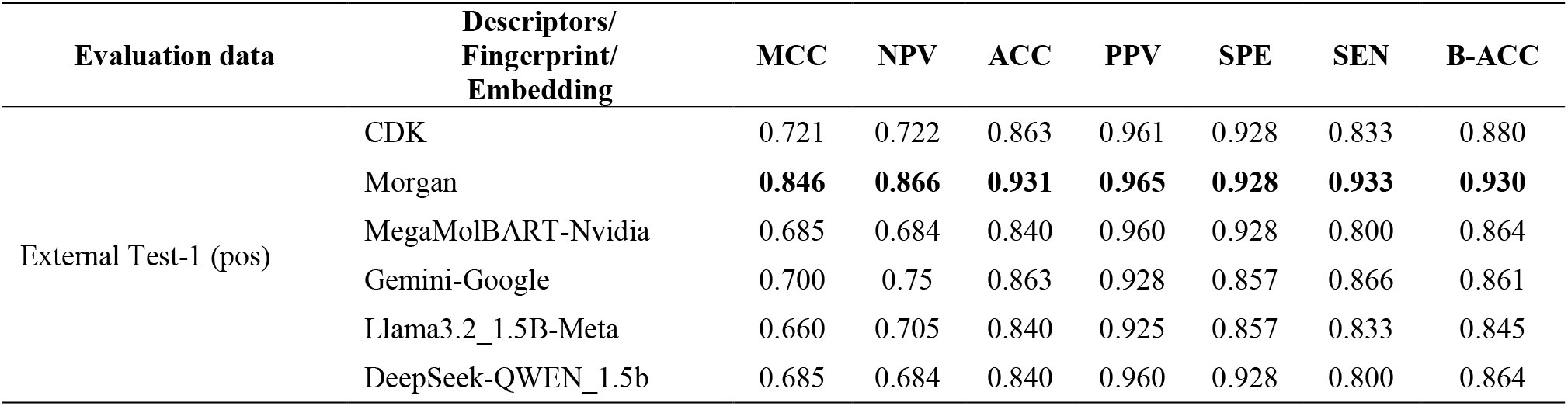
hERG-LTN model performance based on different features input using UnihERG_DB training dataset.

**Table 4:**
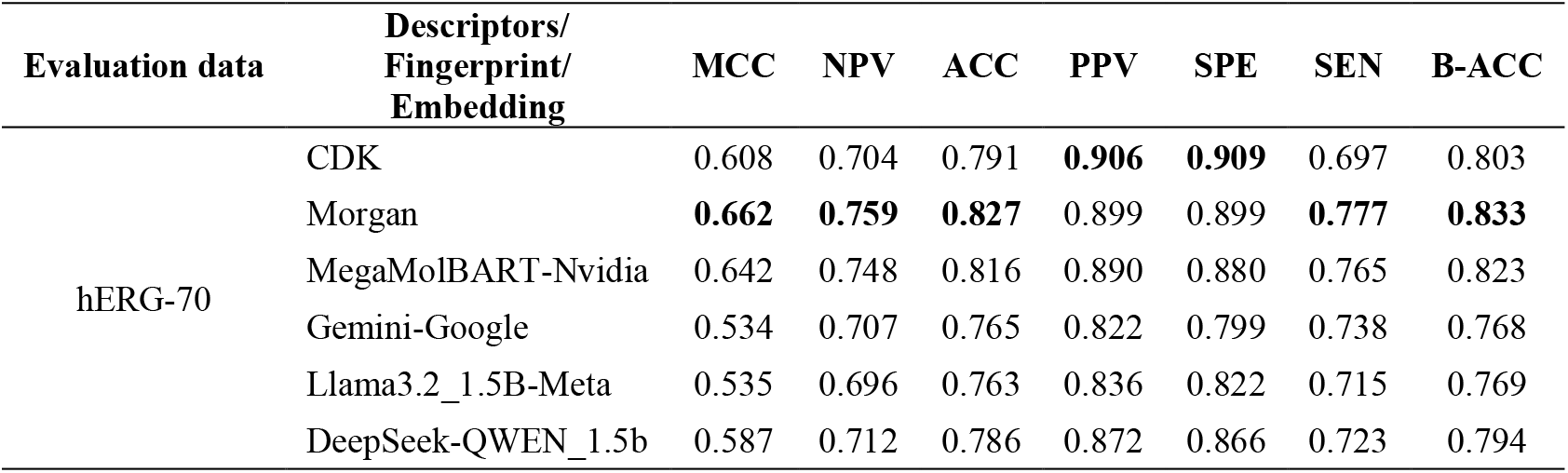
LTN model performance evaluation based on Arab et al. hERG-70.

**Table 5:**
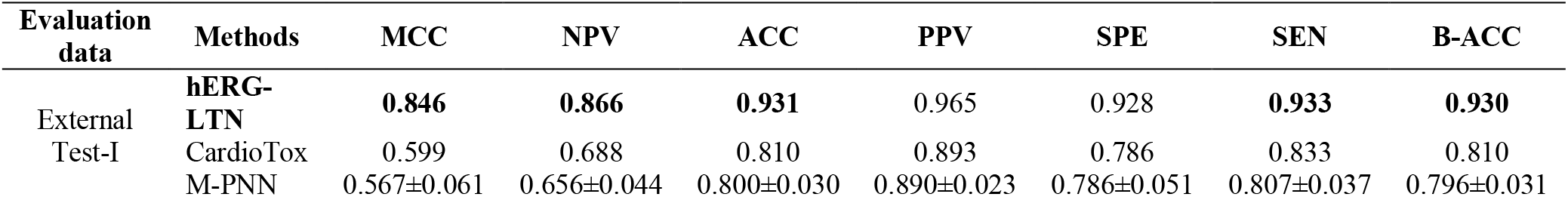
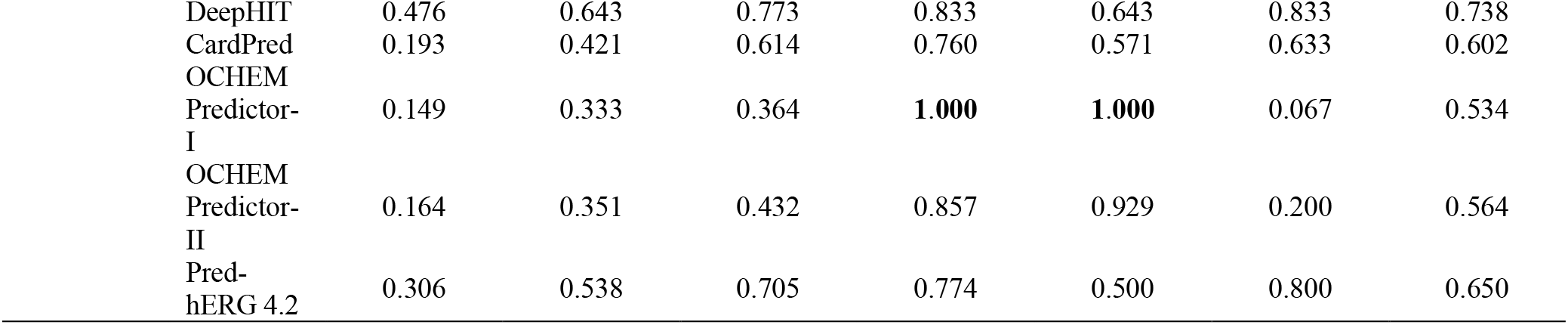
Performance comparison of hERG-LTN classifier on the External Test-1 using UnihERG_DB training data.

**Table 6:** Performance comparison of the hERG-LTN classifier on the hERG-70 benchmark.

| Evaluation data | Methods | ACC | SEN | SPE | F1 | CCR | MCC |
| --- | --- | --- | --- | --- | --- | --- | --- |
| hERG-70 | CardioGenAI | **0.835** | **0.862** | 0.803 | **0.851** | 0.832 | **0.667** |
|  | **hERG-LTN (ours)** | 0.827 | 0.777 | **0.890** | 0.833 | **0.833** | 0.662 |
|  | CToxPred-hERG | 0.814 | 0.867 | 0.746 | 0.839 | 0.807 | 0.621 |
|  | CardioTox | 0.812 | 0.830 | 0.789 | 0.831 | 0.810 | 0.619 |
|  | ADMET1ab 2.0 | 0.717 | 0.716 | 0.718 | 0.738 | 0.717 | 0.431 |
|  | ADMETsar 2.0 | 0.685 | 0.845 | 0.483 | 0.750 | 0.664 | 0.355 |
|  | CardPred | 0.561 | 0.527 | 0.603 | 0.570 | 0.565 | 0.130 |

Table 3 illustrates the hERG-LTN model’s performance based on the External Test-1 dataset. Among the six different types of inputs (CDK, Morgan, MegaMolBART, Gemini-Google, Llama3.2_1.5B-Meta, DeepSeek-QWEN_1.5b), the Morgan features yielded pioneering results, achieving 0.846 (MCC), 0.866 (NPV), 0.931 (ACC), 0.965 (PPV), 0.928 (SPE), 0.933 (SEN), and 0.930 (B-ACC). CDK fingerprints alone are also well competitor with high scores across metrics of 0.721 (MCC), 0.722 (NPV), 0.863 (ACC), 0.961 (PPV), 0.928 (SPE), 0.833 (SEN), and 0.880 (B-ACC). In addition, the experiments incorporated with LLM embeddings (MegaMolBART, Gemini, Llama3.2, DeepSeek-QWEN) demonstrated suboptimal performance compared to Morgan FF. Although Gemini scored higher than other embedding features.

In addition, we evaluated model performance with hERG-70 benchmark data presented by Arab et al. [21] (Table 4). The results indicated that the hERG-LTN model performed well with the Morgan features, achieving an MCC of 0.662, NPV of 0.759, ACC of 0.827, SEN of 0.777, and B-ACC of 0.833. The MegaMolBART-Nvidia embeddings also showed promising results, with an ACC of 0.816, while CDK and Combination were slightly lower. Overall, the results indicate that while all feature sets are practical, Morgan fingerprints provided the most robust performance on this dataset which also depicts in Fig., 2 ROC AUC curve.

**Fig. 2:**
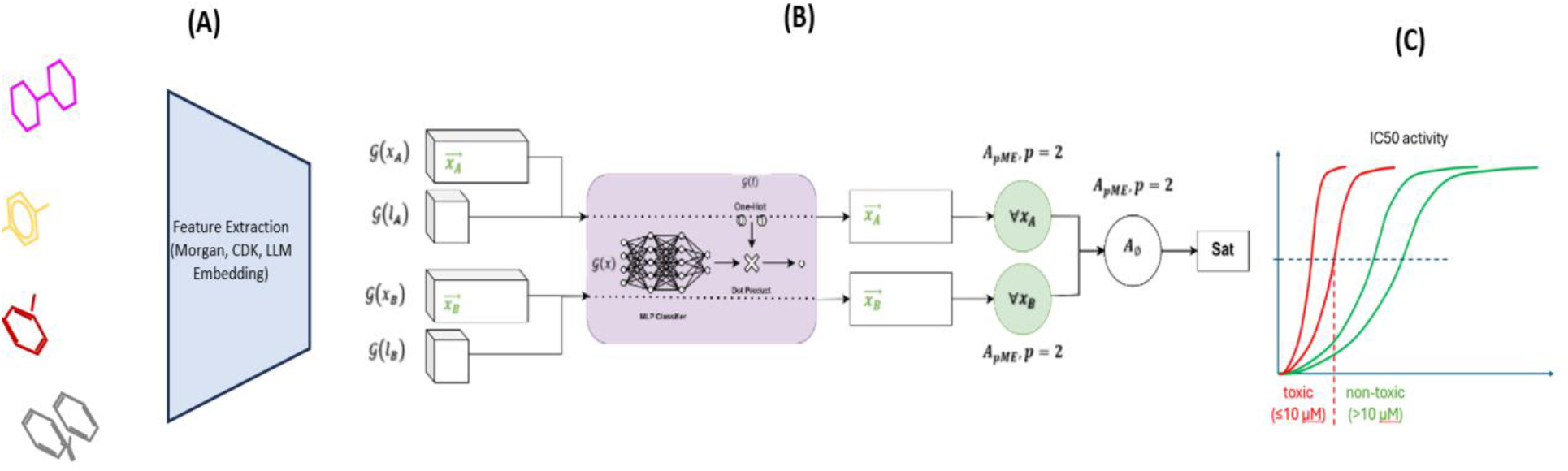
The Binary Classification Architecture of Logic Tensor Network (A)[51]

### A. Benchmarking on External Test-1 and hERG-70 dataset

Further, we compare the performance of our hERG-LTN best classification models to the State-of-the-art models as exhibited in the scientific literature [22]-[25],[28],[1]. We assess with the two different benchmarking External Test-1 (pos) [22]and hERG-70 [21]. Table 5 shows that the hERG-LTN classifier performed exceptionally with morgan features on the External Test-1 (pos), with an 0.846 (MCC), 0.866 (NPV), 0.931 (ACC), 0.965 (PPV), 0.928 (SPE), 0.933 (SEN), and 0.930 (B-ACC). CardioTox was the second-best performer after MCC of 0.599, NPV of 0.688, ACC of 0.810, PPV of 0.893, SPE of 0.786, SEN of 0.833, and B-ACC of 0. 810.OCHEM Predictor-I achieved a PPV of 100% and SPE of 100%.

On the other hand, the hERG-70 benchmark, CardioGenAI (preprint as of Feb 8th) [30], placed the highest ranked, achieving with an ACC of 0.835, SEN of 0.862, F1 score of 0.851, and MCC of 0.667, except SPE, and CCR (Table 6). Though, our hERG-LTN classifier yielded the best scores of SPE (0.890) and CCR (0.833), and other metrics closely followed SOAT models with an ACC of 0.827, SEN of 0.777, SPE of 0.890, F1 score of 0.833, CCR of 0.833, and MCC of 0.662. Alongside, CToxPred-hERG ranked third with an ACC of 0.814, F1 of 0.839, SPE of CCR of 0.807, and MCC of 0.621. Figure. 3 illustrates a Comparison of performance evaluation metric analysis on the hERG-70 dataset.

**Fig 3:**
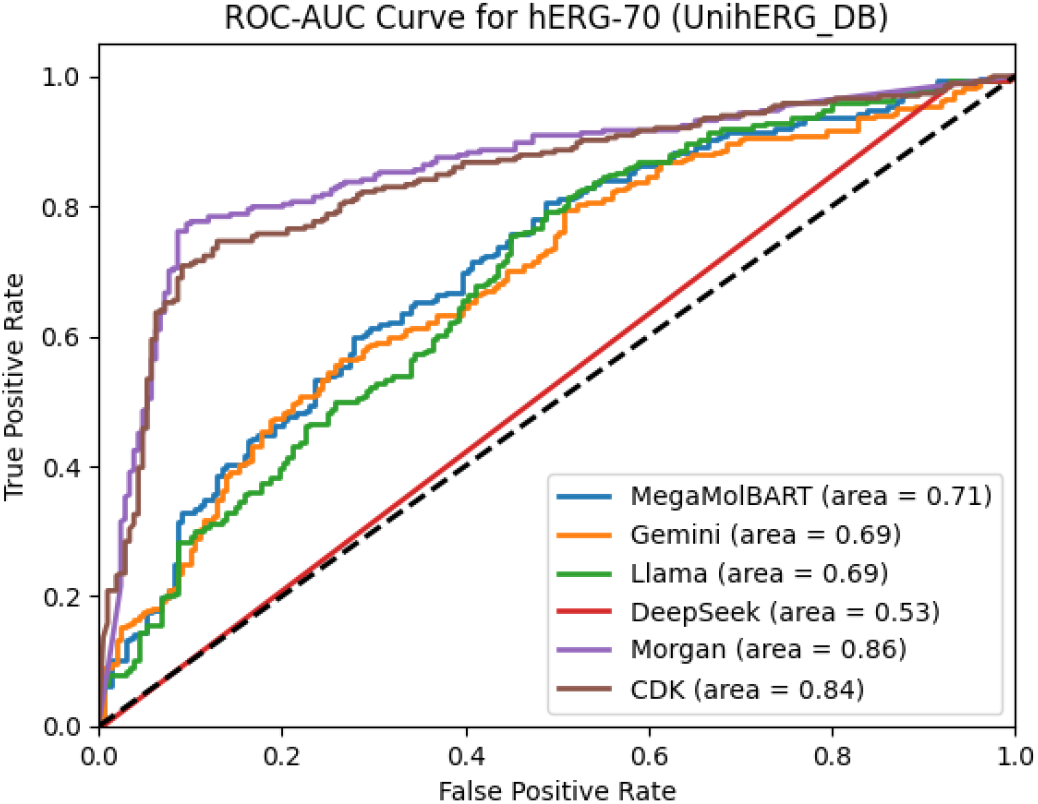
hERG-LTN Model ROC-AUC Curve

## IV. Discussion

A significant challenge in Cardiotoxicity and AI research is the lack of integrated data-knowledge experiments that enable reasoning capabilities. Additionally, current studies are inadequate in leveraging chemical LLM for predicting Adverse drugs. Our findings demonstrate that the hERG-LTN, a Neuro-symbolic approach classifier trained with the UnihERG_DB dataset, is highly effective in predicting hERG-related Cardiotoxicity, significantly outperforming existing models such as CardioTox, M-PNN, and DeepHIT in terms of External test-1 performance (Table 5). In contrast, the hERG-LTN classifier trained on the UnihERG_DB dataset delivers exceptional results, ranking just behind CardioGenAI in the hERG-70 benchmark across ACC, SPE, F1, CCR, and MCC metrics (Table 6). Furthermore, the ROC-AUC curve reveals that the models (CDK, Morgan, MegaMolBART, Gemini, Llama and DeepSeek-QWEN) all show strong performance in predicting hERG-70 inhibition, with AUC values ranging from 0.84 to 0.86.

The study has some limitations that should be considered. The model’s interpretability and explainability are limited as it is a complex neural network, and further work is needed to improve its transparency. The model’s performance on the hERG-70 benchmark is slightly lower than some state-of-the-art models, indicating room for improvement.

## V. Conclusions

In conclusion, the article emphasizes that the Neuro-Symbolic AI approach employed in the hERG-LTN classifier establishes a new paradigm for hERG-related cardiotoxicity evaluation, markedly surpassing existing models. This study explored various methodologies, including molecular fingerprinting, descriptors, and LLM embeddings. The synthesis of data and knowledge-driven strategies reveals immense potential for propelling precision pharmacogenomics in drug discovery and assessing drug-induced Cardiotoxicity. Future endeavors could involve experimenting with other avantgarde Generative A.I. models, aggregating more heterogeneous datasets, and further investigating alternative Neuro-Symbolic AI frameworks to augment predictive accuracy and model resilience in hERG cardiotoxicity research.

## Acknowledgment

We extend our deepest gratitude to SPARC and acknowledge that this project was generously funded through grant OT2:lOT20D032742-Ol and UMI TR004771, whose support made this study possible.

